# Bystander base editing interferes with visual function restoration in Leber congenital amaurosis

**DOI:** 10.1101/2024.10.23.619839

**Authors:** Seok-Hoon Lee, Jun Wu, Dongjoon Im, Gue-ho Hwang, You Kyeong Jeong, Hui Jiang, Seok Jae Lee, Dong Hyun Jo, William A. Goddard, Jeong Hun Kim, Sangsu Bae

## Abstract

Base editors (BEs) have emerged as a powerful tool for gene correction with high activity. However, bystander base editing, a byproduct of BEs, presents challenges for precise editing. Here, we investigated the effects of bystander edits on phenotypic restoration in the context of Leber congenital amaurosis (LCA), a hereditary retinal disorder, as a therapeutic model. We observed that in *rd12* of LCA model mice, the highest editing activity version of an adenine base editors (ABEs), ABE8e, generated substantial bystander editing, resulting in missense mutations despite RPE65 expression, preventing restoration of visual function. Through AlphaFold-based mutational scanning and molecular dynamics simulations, we identified that the ABE8e-driven L43P mutation disrupts RPE65 structure and function. Our findings underscore the need for more stringent requirements in developing precise BEs for future clinical applications.

## Introduction

The CRISPR-Cas system is a powerful tool for gene disruption with high efficacy(Jinek *et al*, 2012). CRISPR-Cas9 nucleases generate DNA double strand breaks (DSBs) at target sites in a single-guide RNA (sgRNA)-dependent manner, after which cleaved DNA is repaired by cellular repair pathways, frequently resulting in gene disruption. Owing to this advantage, the first CRISPR drug, named Exa-Cel (Casgevy) was approved by the Food and Drug Administration of the United Kingdom and United States in 2023. This drug disrupts hemoglobin subunit beta-related gene for treating transfusion-dependent beta thalassemia and severe sickle cell disease(Frangoul *et al*, 2021). However, such gene disruption strategies may not be applicable for other genetic diseases, which may require gene correction, including base correction, rather than gene disruption. Moreover, CRISPR-Cas nuclease-driven DSBs can cause large deletions, chromosomal depletions, translocations, P53-mediated cell death, and cellular senescence, potentially hindering therapeutic applications.

Therefore, base editors (BEs) have attracted great attention as a gene correction drug because they can convert one or a few substitutions with high editing efficacy without creating DNA DSBs. BEs mainly consist of a partially deactivated Cas protein, such as the Cas nickase, which is essential for target recognition and unwinding of the DNA duplex, and a specific deaminase that catalyzes nucleotide conversion. Various BE platforms have been developed, which involve a cytosine BE (CBE) for C-to-T conversion(Komor *et al*, 2016), an adenine BE (ABE) for A-to-G conversion(Gaudelli *et al*, 2018), a cytosine transversion BE (CGBE1) for C-to-G conversion(Kurt *et al*, 2021), and an adenine transversion BE (AYBE) for A-to-T and A-to-C conversion(Tong *et al*, 2023). Thus, it is possible to correct all types of base substitutions by selecting appropriate BE platforms.

However, BEs have several undesired limitations as follows: i) sgRNA-dependent off-target edits in the genome, which can be addressed by using high-fidelity Cas proteins(Hu *et al*, 2018; Lee *et al*, 2018; Rees *et al*, 2017), ii) sgRNA-independent off-target edits in DNA or RNA, which can be mitigated by engineering deaminases(Grünewald *et al*, 2019; Rees *et al*, 2019; Zhou *et al*, 2019), and iii) bystander edits at on-target sites within editing activity windows, which might interfere with functional restoration(Jeong *et al*, 2020). Among them, it is particularly challenging to completely avoid bystander edits, although a few studies have suggested BE variants with narrower editing windows(Kim *et al*, 2017; Liu *et al*, 2020). Furthermore, there is a trade-off between editing efficacy and specificity, with BE variants that have higher editing efficiency showing higher bystander editing rates. However, the functional effect of bystander edit-driven missense mutations has not been comprehensively validated at the animal level.

Leber congenital amaurosis (LCA) is a representative inherited retinal disorder causing blindness in childhood. RPE65, a gene responsible for converting all-*trans*-retinyl esters to 11-*cis*-retinol, is one of the major factors triggering LCA(Kiser, 2022). Most patients with pathogenic mutations in *Rpe65* have severe visual impairment during childhood and adolescence. We and other groups have tried to rescue retinal degeneration 12 (*rd12*) model mice that harbor a nonsense mutation in *Rpe65* (c.130C>T, p.R44X), identical to the mutation causing LCA in the Chinese population(Pang *et al*, 2005). Palczewski et al. first used ABEmax by a lentiviral delivery method(Suh *et al*, 2021) and our group utilized NG-ABEmax (TadAmax based on NG protospacer-adjacent motif [PAM]-targetable SpCas9) with adeno-associated virus (AAV) split delivery(Jo *et al*, 2023). In addition, prime editors were used to achieve more precise treatment of *rd12* mice(Jang *et al*, 2022).

Overall, ABEs could edit the pathogenic mutation in *Rpe65* with high efficacy, resulting in the restoration of visual function in *rd12* mice. The induction of undesired bystander editing near the target mutation by ABEs has not been studied in detail. In this study, we compared ABE variants for treating *rd12* model mice and comprehensively evaluated the genotype and phenotype of undesired bystander editing effects. We examined the relationship between bystander edits and functional recovery using AlphaFold-based mutational scanning and molecular dynamics (MD) simulations. These findings highlight the importance of understanding the effects of BE-mediated bystander editing in the development of gene correction therapies.

## Results

### Different DNA editing outcomes generated by the three ABE variants

The *rd12* mouse model contains a homozygous nonsense mutation in *Rpe65* (c.130C>T, p.R44X), making it a representative LCA model(Pang *et al*., 2005). The target adenine (A6) causing the nonsense mutation can be corrected by ABE variants, but several bystander nucleotides around it, including A3, A8, A11, and C5 with a TC motif, can be targeted by ABEs (Fig. 1A)(Jo *et al*., 2023). To compare and identify ABE variants with high on-target editing but low bystander editing activities, we employed three representative ABE variants based on NG-Cas9; i) ABEmax, the first optimized version created by Liu et al., ii) ABE8e, known for its high editing activity, and iii) ABE8eWQ, which our group reported to have minimal bystander TC edits and transcriptome-wide RNA deamination effects. ABE8e has a wider editing window (positions 3–11, counting the end distal to the PAM sequence as position 1 and higher editing efficiency compared with ABEmax (positions 4–8) and ABE8eWQ (positions 4–8)(Jeong *et al*, 2021; Richter *et al*, 2020).

**Figure 1.**
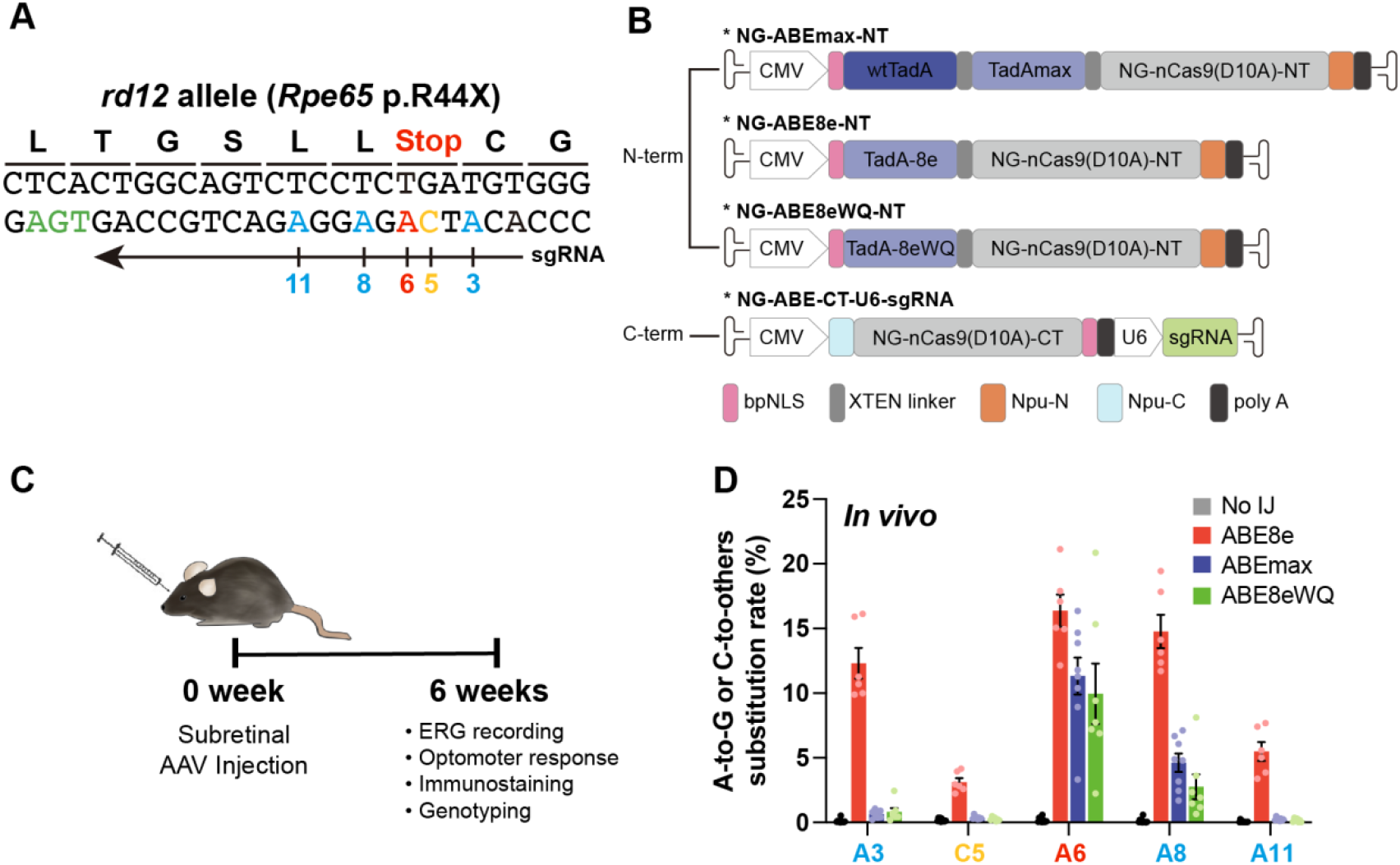
Different DNA editing outcomes generated by the three ABE variants. **(A)** DNA context around the nonsense mutation of *rd12* mice. The arrow indicates the sgRNA for NG-ABEs and colors indicate target adenosine (red), bystander adenosine (blue), bystander cytosine (yellow), and PAM (green). Nucleotide number indicates position, counting PAM as position 21–23. **(B)** Schematic drawing of the dual-AAV vectors for ABE delivery. CMV and U6 are promoters. Npu-C and Npu-N indicate C- and N-intein from *N. punctiforme*, respectively. **(C)** Schematic showing outline of *in vivo* experiments. **(D)** High-throughput sequencing results of the nonsense mutation region in the genomic DNA isolated from RPE tissue of no injection (No IJ) (n = 11), ABE8e-treated (n = 6), ABEmax-treated (n = 8), and ABE8eWQ-treated mice (n = 7). The split-AAV strategy was utilized to deliver ABEs, and each component of split ABEs was packaged into AAV2/9. Error bars indicate mean ± s.e.m.

For *in vivo* delivery of ABE variants into *rd12* mouse, all ABE variants were prepared using dual adeno-associated virus (AAV) vectors in a split form with a trans-splicing intein, due to the limited size capacity of AAVs. One vector contained the N-terminal part of ABEs (TadAmax, TadA-8e, and TadA-8eWQ with the N-terminal part of NG-Cas9 nickase), whereas the other vector contained the C-terminal part of the ABE (C-terminal part of NG-Cas9 nickase) along with sgRNA (Fig. 1B). Prior to *in vivo* injection experiments, we compared the editing outcomes of ABEmax, ABE8e, and ABE8eWQ in mouse embryonic fibroblasts (MEFs) from *rd12* mice. In *rd12* MEFs, ABE8e exhibited the highest editing efficiency both at the target A6 (7.22%) and at bystanders (4.55% of A3, 0.46% of C5, 6.47% of A8, and 0.32% of A11). By contrast, ABEmax and ABE8eWQ showed relatively lower editing efficiency at target A6 (5.23% and 2.67%, respectively) and bystander A8 (0.98% and 0.18%, respectively) (Fig. EV1). Notably, both ABEmax and ABE8eWQ exhibited negligible editing efficiency at bystanders of A3, C5, and A11.

Next, three dual AAVs (serotype AAV2/9) for ABEmax, ABE8e, and ABE8eWQ were constructed and injected into the subretinal region of 3-week-old *rd12* mice. After 6 weeks, functional recovery, RPE65 levels, and genotyping were evaluated (Fig. 1C). High-throughput sequencing analysis of genomic DNA from ABE-injected *rd12* mice revealed that ABE8e had the highest editing efficiency both at the target A6 (average 16.38%, n = 6) and at bystanders (average 12.32% at A3, 3.12% at C5, 14.77% at A8, and 5.49% at A11, n = 6). By contrast, ABEmax and ABE8eWQ showed relatively lower editing efficiency at target A6 (average 11.33% and 9.96%, n = 8 and n = 7, respectively) and bystander A8 (average 4.61% and 2.76%, n = 8 and n = 7, respectively) with negligible editing efficiency at other bystander sites A3, C5, and A11. Consequently, ABEmax showed a similar editing frequency to ABE8eWQ in RPE tissue (Fig. 1D). These results are very similar to those obtained in *rd12* MEFs (Fig. EV1).

Overall, all bystander adenines and cytosine were highly converted by ABE8e, whereas only bystander A8 was converted by ABEmax and ABE8eWQ. Therefore, the resulting missense mutations are not critical. We expected that ABE8e would exhibit the highest level of visual restoration because ABE8e resolved the premature stop codon more efficiently than other ABEs.

### Rescue of RPE65 expression could not restore visual function in *rd12* mice

To evaluate changes in molecular and visual function in ABE-treated *rd12* mice, we used C57BL/6 mice as positive controls and ABE-untreated *rd12* mice as negative controls. Six weeks after ABE injection, the retinal pigment epithelial–choroid–sclera (RCS) complex was dissociated and processed as a wholemount. Immunofluorescence staining showed RPE65 expression in RPE tissue from C57BL/6 and ABE-treated mice, but not untreated mice (Fig. 2A). The percentage of RPE65-positive cells was counted in randomly selected immunostaining fields, showing recovery rates of 53.7%, 50.8%, and 49.7% in ABE8e-, ABEmax-, and ABE8eWQ-treated mice, respectively (Fig. 2B). These data correlated well with the base correction frequencies at on-target A6 (Fig. 1D), indicating that the premature stop codon was resolved and full RPE65 was produced.

**Figure 2.**
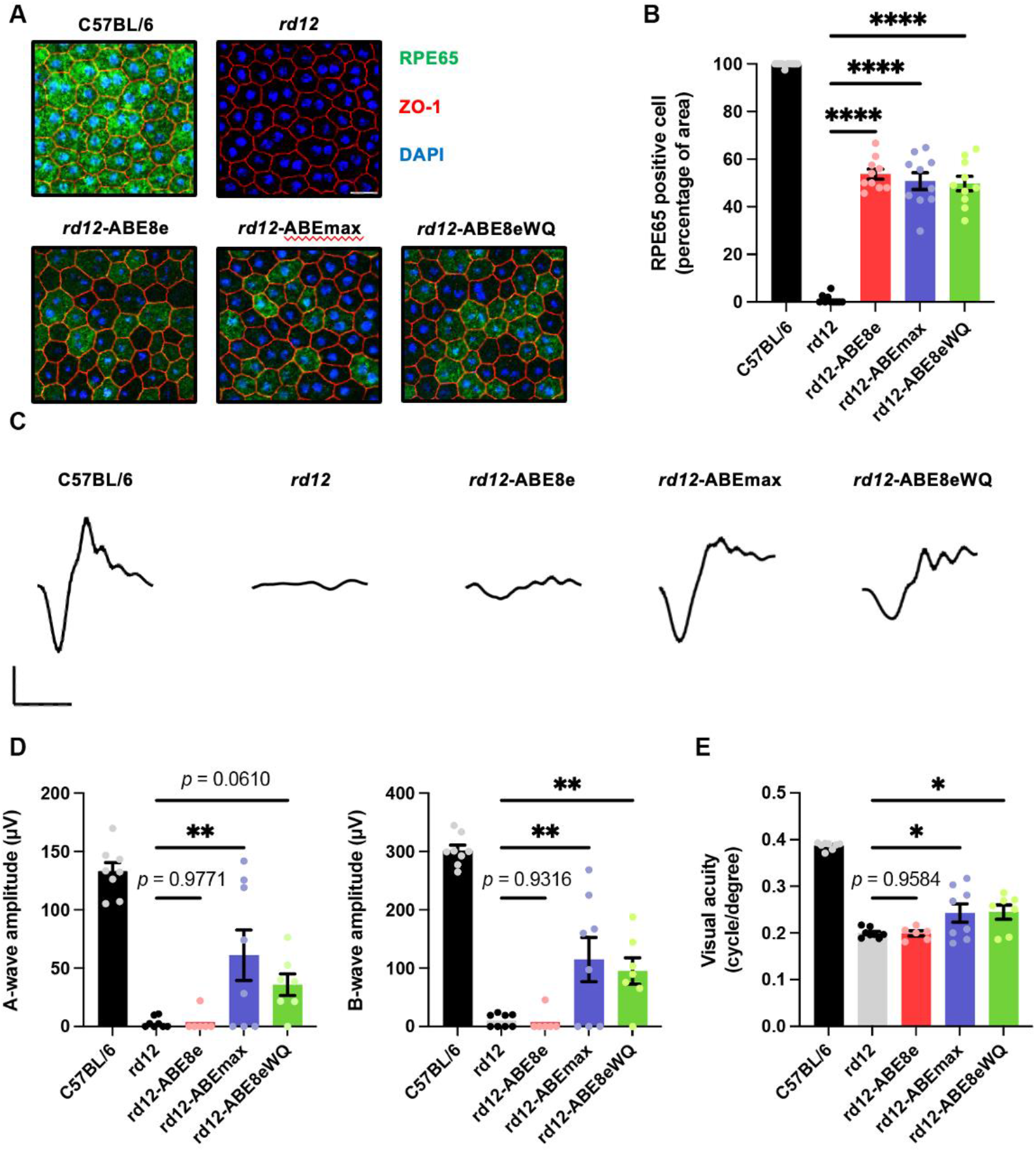
Rescue of RPE65 expression cannot restore visual function effectively in *rd12* mice. **(A)** Representative confocal photomicrographs after immunostaining to show RPE65 expression in RPE cells of C57BL/6, *rd12*, and ABE-treated *rd12* mice. Scale bars: 20 μm. Green indicates RPE65, red indicates ZO-1 (marker for tight junctions), and blue indicates DAPI staining. **(B)** Quantification of RPE65-positive cells from RCS wholemount (n = 10). **(C)** Representative scotopic ERG waveforms from C57BL/6, *rd12*, and ABE*-*treated *rd12* mice. Scale bars: 30 ms (*x*-axis), 100 μV (*y*-axis). **(D)** Quantitative analysis of amplitudes of a- and b-waves of scotopic response (n = 8 for C57BL/6, *rd12*, and ABEmax-treated mice, n = 6 for AAV8e-treated mice, n = 7 for AAV8eWQ-treated mice). **(E)** Quantitative analysis of the visual acuity of C57BL/6, *rd12*, and ABE*-*treated *rd12* mice (n = 8 for C57BL/6, *rd12*, and ABEmax-treated mice, n = 6 for AAV8e-treated mice, n = 7 for AAV8eWQ-treated mice). \**p* < 0.05, \*\**p* < 0.01, \*\*\*\**p* < 0.0001. Error bars indicate mean ± s.e.m.

Next, visual chromophore recovery was determined by electroretinography (ERG) for ABE-treated mice. Contrary to the RPE65 expression data, the ERG waveforms were not recovered in some ABE-treated mice and exhibited an opposite tendency. Notably, in ABE8e-treated mice, the amplitude of a- and b-waves of scotopic responses was on average, 2.7% and 2.5% that of wild-type mice, respectively. By contrast, definite responses to bright stimuli were observed in ABEmax- and ABE8eWQ-treated mice, which were significantly higher than that in untreated mice and ABE8e-treated mice (Fig. 2C and 2D). Optomotor responses to rotating stimuli in a virtual cylinder were measured. Significant recovery of visual thresholds was detected in ABEmax- and ABE8eWQ-treated mice, whereas no significant difference was observed in ABE8e-treated mice compared with untreated mice (Fig. 2E). The inconsistency in ABE8e results between sequencing and immunofluorescence staining data versus visual function restoration might be caused by undesired bystander editing or other off-target editing of the genome and RNA transcripts.

### Comprehensive identification of undesired editing outcomes induced by ABEs in ABE-treated *rd12* mouse

We sought to determine why ABE8e-treated mice exhibited worse visual function recovery despite higher editing efficiency than ABEmax- and ABE8eWQ-treated mice. To this end, we first investigated possible sgRNA-dependent off-target editing using Cas-OFFinder software. Allowing for up to two mismatched bases, we identified 16 potential off-target sites: 6 with mismatches within the PAM-distal half (OT1-OT6) and 10 with mismatches within the PAM-proximal half (OT7-OT16) (Fig. EV2A). High-throughput sequencing revealed off-target editing at three sites (OT1, OT3, and OT4) only in ABE8e-treated mice (Fig. 3A and Fig. EV2B). However, because these off-target sites were in non-coding regions, they were unlikely to affect RPE65 production and visual recovery. Next, we investigated sgRNA-independent off-target edits on RNA transcripts by measuring A-to-I conversion frequencies in three representative RNA transcripts (*AARS1, MCM3AP*, and *PERP*). High-throughput sequencing results showed mild RNA edits in ABE8e- and ABEmax-treated mice, but none in ABE8eWQ-treated mice (Fig. 3B). These results suggest that RNA off-target edits are not the primary reason for impaired visual restoration because although ABEmax also showed mild RNA off-target edits, it exhibited moderate functional recovery (Fig. 2C–2E). Last, we focused on bystander editing effects. The results represented bulk conversion rates of each base (Fig. 1B and 1D). We examined every pattern of editing outcomes in mice treated with the three ABE variants and found that 50% of the editing outcomes were intended RPE65 patterns (called “precise”) with ABEmax and ABE8eWQ (average 6.40% and 6.90%, respectively), whereas only a small portion (average 1.33%) of editing outcomes was precise among all editing outcomes (average 16.30%) with ABE8e in RPE tissue (Fig. 3C). Taken together, bystander editing plays a major role in inhibiting visual function recovery by ABE8e.

**Figure 3.**
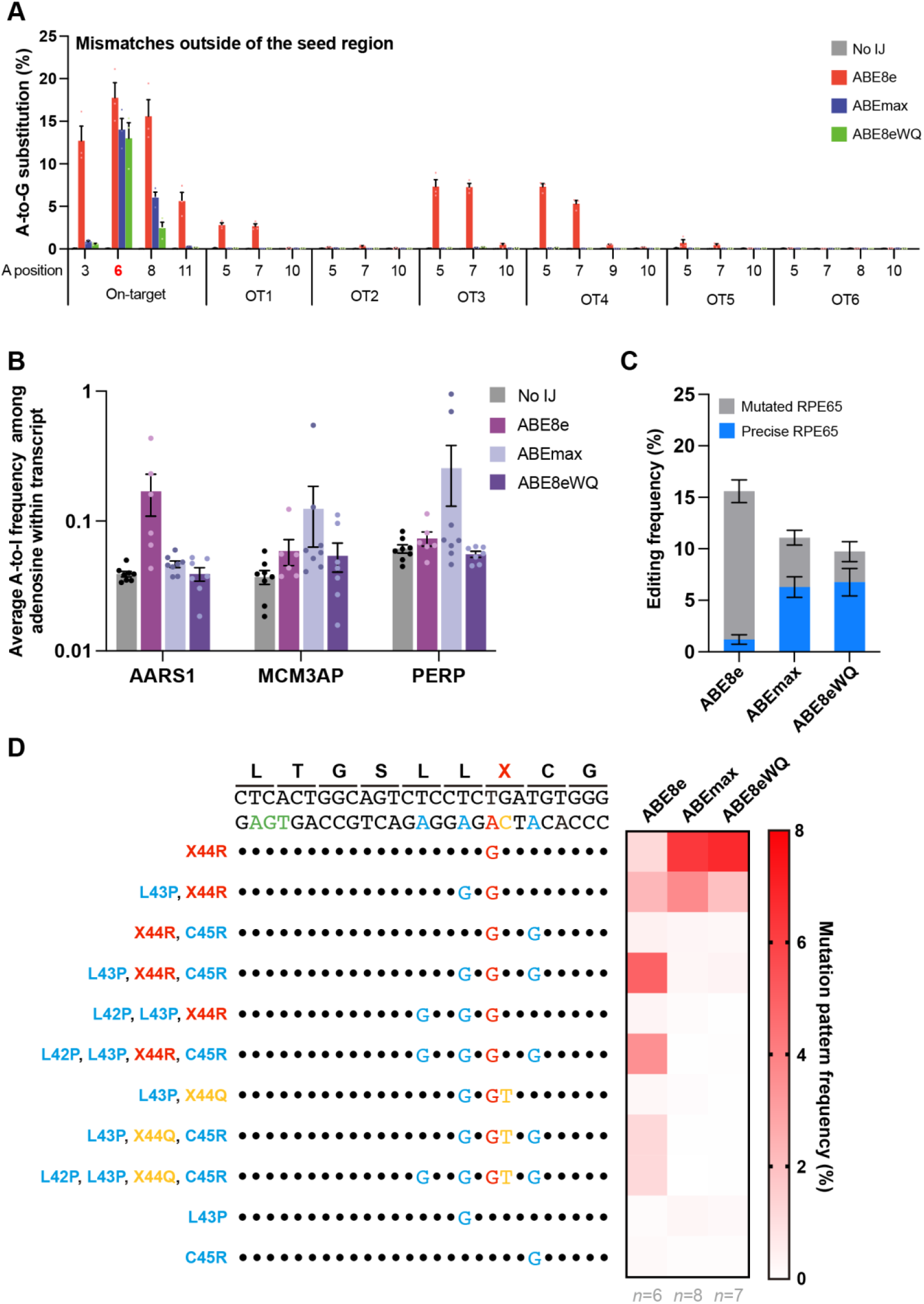
Comprehensive identification of undesired editing outcomes induced by ABEs in ABE-treated *rd12* mouse. **(A)** Frequencies of sgRNA-dependent off-target edits in genomic DNA isolated from the RPE tissue of ABE-treated mice (n = 3). The positions of adenosine on each off-target site are described below the *x*-axis. OT1-OT6 are sgRNA-dependent off-target sites that contain mismatches outside the sgRNA seed region. **(B)** A-to-I conversion frequencies in the three mRNA transcripts after treatment with the three ABE variants. RNA was extracted from the RPE tissue of no injection (No IJ) (n = 8), ABE8e-treated (n = 6), ABEmax-treated (n = 8), and ABE8eWQ-treated mice (n = 7). **(C)** Frequencies of precise RPE65 and mutated RPE65 in genomic DNA isolated from the RPE tissue of ABE-treated mice (n = 6 for ABE8e, n = 8 for ABEmax, and n = 7 for ABE8eWQ). **(D)** Average mutation pattern frequencies with the three ABE variants in genomic DNA isolated from RPE or retina tissue. Amino acid substitutions are listed on the left of each DNA mutation pattern. Colors indicate target adenosine (red), bystander adenosine (blue), bystander cytosine (yellow), and PAM (green). Black dot indicates the same nucleotide with reference. Error bars indicate mean ± s.e.m.

We analyzed substitution rates based on major editing patterns. Numerous mutated RPE65 variants were generated by ABE because each bystander A or C can trigger missense mutations of different amino acids. In ABE8e, RPE65 variants containing L43P, C45R or L42P, L43P, and C45R accounted for the highest proportion (4.96% and 3.51% from RPE tissue, respectively). ABE8e showed significant bystander C editing, generating RPE65 variants containing L43P, R44Q, C45R or L42P, L43P, R44Q, and C45R (1.21% and 1.19% from RPE tissue, respectively). Bystander C editing disrupts the correctly edited X44R as R44Q. By contrast, precise RPE65 was the most frequent outcome when *rd12* mice were injected with ABEmax or ABE8eWQ (6.28% and 6.75% from RPE tissue, respectively). However, ABEmax and ABE8eWQ could not completely avoid bystander effects. The L43P RPE65 mutant was also generated by ABEmax and ABE8eWQ at high frequencies (3.68% and 1.98% from RPE tissue, respectively) (Fig. 3D). Overall, of all the undesired editing outcomes, we hypothesize that bystander editing causes insufficient visual restoration.

### Detailed analysis of bystander effect in the *rd12* mouse model and at the molecular level of RPE65

Each ABE-treated *rd12* mouse exhibited different editing efficiency and levels of visual restoration, even when using the same version of ABE, due to variations in ABE delivery efficiency and interindividual differences between mice. Therefore, we collected all *in vivo* genotyping and phenotyping data regardless of the ABE version, to investigate whether missense mutations interfere with visual restoration. We arranged ABE-treated mice according to the degree of visual recovery, along with the distribution of correct or mutated RPE65. The top 10 mice showed a large proportion of precise RPE65, whereas the bottom 10 mice exhibited a small proportion of precise RPE65 and a relatively large portion of mutated RPE65 (Fig. 4A). Additionally, we compiled all *in vivo* genotypic and phenotypic data from ABE-treated *rd12* mice, regardless of the ABE version, and examined the correlation between RPE65 mutation patterns and visual restoration (b-wave amplitude or visual acuity). Through correlation analysis, we verified that overall bystander editing interferes with visual restoration (Fig. 4B and Fig. EV3). All mutated RPE65 harbor the L43P mutation, leading us to hypothesize that L43P is the primary factor hindering visual restoration.

**Figure 4.**
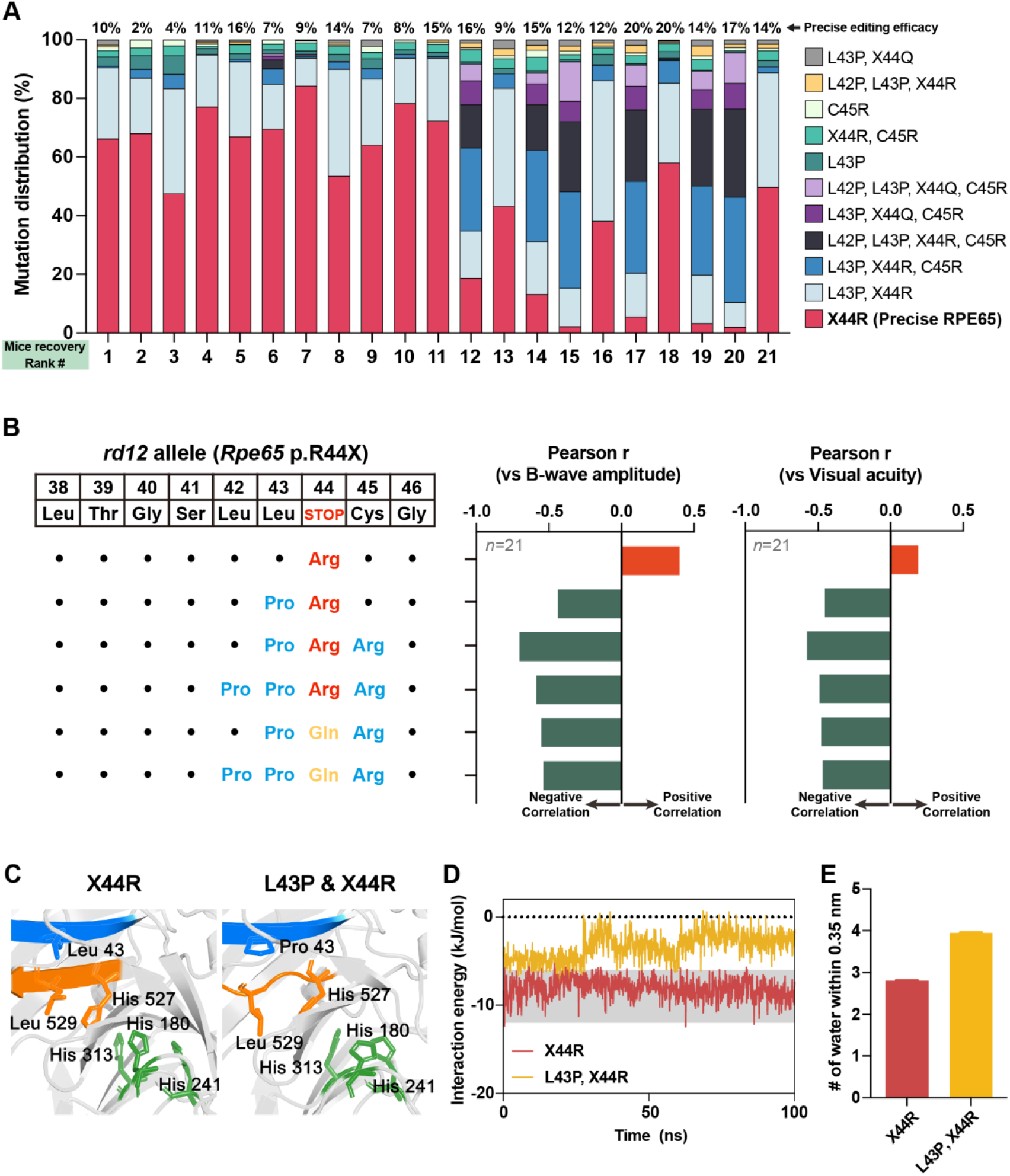
Detailed analysis of the bystander effect in the *rd12* mouse model and at the molecular level of RPE65. **(A)** The distribution of precise or mutated RPE65 in each mouse in accordance with the sequence of successful recovery (n = 21; collected from *rd12* mouse treated with the three ABE variants). Precise editing efficacies are described on the upper side of each column. X44R indicates precise RPE65. **(B)** Correlation between mutation pattern of RPE65 and two phenotypes representing visual recovery (ERG B-wave amplitude and visual acuity). Black dot indicates the same amino acid with reference. Orange and green columns indicate positive and negative Pearson’s correlation, respectively. **(C)** Comparison of the equilibrated MD structure between precise and L43P RPE65. The ^42^LLRC^45^ and ^527^HGLF^530^ domains are shown in blue and orange, respectively, and the mutated or affected side chains (i.e., Leu43, His180, His241, His313, His527, and Leu529) are depicted as sticks. **(D)** Simulated interaction energies for carbon backbone atoms of the ^42^LLRC^45^ and ^527^HGLF^530^ domains of normal and L43P RPE 65. **(E)** Average number of water molecules within 0.35 nm around the ^42^LLRC^45^ domain during MD simulation. Error bars indicate mean ± s.e.m.

Consistent with our *in vivo* results, the function of RPE65 was impaired by L43P, in HEK293 cells(Suh *et al*., 2021). However, contrary to our results in *rd12* mice, the L43P mutation lowered the stability of RPE65 in HEK293 cells, based on immunoblot analysis. We believe that the effects of bystander editing on the structure of RPE65 must be investigated further because our *in vivo* immunostaining results demonstrated that the recovered RPE65 structure can bind antibodies (Fig. 2A). Therefore, we performed AlphaFold-based mutational scanning to understand the negative effects of bystander editing on visual restoration (Fig. EV4). To systematically examine the impact of point mutations on the ^42^LLRC^45^ domain, which is the on-target site of ABE, we employed AlphaFold to predict the structure of RPE65 and its variants (Fig. EV4). The ^42^LLRC^45^ and ^527^HGLF^530^ domains of RPE65 normally stack with beta sheet structures (Fig. EV4). However, the L43P point mutation disrupted the beta sheet structure of the ^527^HGLF^530^ domain, but not the ^42^LLRC^45^ domain itself (Fig. 4C and Fig. EV5). This correlation with our experiments indicates that the L43P missense mutation from bystander editing induces a negative effect on phenotype recovery *in vivo*.

Based on these predictions from AlphaFold, we investigated atomistic details using MD simulations (see details in Methods section) to observe the interactions between the beta sheet structured domains (^42^LLRC^45^ and ^527^HGLF^530^) and the structural conversion of RPE65 induced by the L43P missense mutation. The AlphaFold-predicted structures were employed as an initial structure for 100 ns MD at 310 K to obtain equilibrated structures for RPE65 and its L43P mutant (Fig. 4C). The global shape of RPE65 and the L43P mutant changed little (RMSD = 1.371 Å), but a dramatic structural rearrangement occurred around the ^42^LLRC^45^ and ^527^HGLF^530^ domains (Fig. 4C). The interaction energy between the carbon backbone atoms of the ^42^LLRC^45^ and ^527^HGLF^530^ domains of wild-type RPE65 did not change significantly throughout the simulation (Fig. 4D). By contrast, the interaction of the initial state for the mutated form was relatively weak and changed after 30 ns with decreased interaction energy (Fig. 4D). Interestingly, the average number of water molecules around the ^42^LLRC^45^ domain increased due to the L43P point mutation, disrupting hydrophobic interactions between the ^42^LLRC^45^ and ^527^HGLF^530^ domains (Fig. 4E and Fig. EV6). Taken together, the initial structure of the L43P RPE65 mutant was relatively unstable with large local structural rearrangements occurring during MD simulation. Our results suggest that the L43P missense mutation, frequently generated through ABE bystander editing, mediates structural changes near the catalytic site and impairs RPE65 function.

### Discussion

Here, we identified the effect of ABE bystander editing triggering missense mutation in *rd12* mice representing LCA. Immunofluorescence staining results showed a high frequency of stop codon release and substantial formation of the RPE65 structure, capable of antibody binding, following the administration of the three ABE variants to *rd12* mice. However, the visual restoration of ABE8e-treated mice was poor, which is in contrast to the results of immunofluorescence staining. We found that the missense mutations generated by bystander editing have a negative correlation with phenotypic restoration. AlphaFold-based mutational scanning and MD calculations revealed that in the ^42^LLRC^45^ *β*-sheet, L43P induced loss of *β*-sheet structure of the ^527^HGLF^530^ sequence, which in turn modified the adjacent catalytic site. To our knowledge, this study is the first to report that bystander editing can interfere with sufficient functional recovery in mice.

Sufficient on-target editing efficacy is required to qualify as a gene editing drug. ABE8e typically has the highest editing efficacy compared with other ABEs(Richter *et al*., 2020). However, along with improved editing efficacy, the editing window is also expanded. This expanded editing window can lead to more bystander editing, triggering missense mutations and resulting in poor phenotypic restoration. Bystander editing is inevitable even with other BEs characterized by narrow editing window or when selecting different sgRNAs to minimize it. Therefore, when applying BEs as CRISPR therapy, targets with a high likelihood of generating missense mutations through bystander editing should be avoided. Alternatively, non-viral delivery of BE ribonucleoprotein complexes should be considered to minimize bystander editing, although some level of bystander editing may still occur. Additionally, missense mutations generated by bystander editing must be examined, at least at the animal or molecular level, through structural prediction and MD simulation.

The PE platform can be a promising alternative to avoid concerns of bystander editing, which comprises a reverse transcriptase fused to an RNA-programmable nickase and a prime editing guide RNA(Anzalone *et al*, 2019). PE can introduce insertions, deletions, and various substitutions without causing DSBs. Recent studies have shown that the use of AAV-PEs restored RPE65 expression and improved visual function in *rd12* mice(Jang *et al*., 2022). However, despite its versatility, the editing efficiency and phenotype recovery of PE were not completely satisfactory compared with BE. Moreover, efforts to package PE with dual-AAV systems have encountered challenges due to its large size(Davis *et al*, 2023; Doman *et al*, 2023), making it difficult to include all necessary components, such as factors that enhance editing efficacy, including hMLH1dn, in dual-AAV vectors. Although PE offers advantages, such as larger targeting scope and reduced undesired editing, over Cas nucleases or BEs, further improvement and optimization are needed for its application as CRISPR therapy. Thus, BEs are considered a more viable option for base correction therapy. In this context, our study demonstrates that bystander editing interferes with phenotypic restoration in animals. These findings underscore the importance of developing more precise BEs for clinical applications.

## Methods

### Molecular cloning and virus production

All plasmids were constructed with the Gibson assembly cloning method. An N-terminal coding sequence of ABEmax in the AAV2-ITR backbone was utilized, which was constructed earlier by our group. To construct the N-terminal part of ABE8e and ABE8eWQ in the AAV2-ITR backbone, the N-terminal part of ABEmax was digested by NotI (NEB, R3189L) and BglII (Enzynomics, R010S). For insert fragment preparation, each TadA region was amplified with Phusion DNA polymerase (Thermo Fisher Scientific, F530L). The digested AAV2-ITR backbone and amplified PCR products were purified with a gel extraction kit (Expin Gel SV mini; GeneAll, 102-102). The digested AAV2-ITR backbone and PCR product were mixed in a volume of 10 μL, containing 2 U of T5 exonuclease (NEB, M0363S), 12.5 U of Phusion DNA polymerase (Thermo Fisher Scientific, F530L), 2 kU of Taq DNA ligase (NEB, M0208S), 0.2 M Tris-HCl (pH 7.5), 0.2 M MgCl_2_, 2 mM dNTPs, 0.2 M dithiothreitol, 25% PEG-8000, and 1 mM NAD, and incubated at 50°C for 1 h. The mix was transformed into 50 μL of C3040 competent cells. A single colony was picked and inoculated into LB medium containing antibiotics. Recombinant AAV packaging (AAV-NT-ABEmax, AAV-NT-ABE8e, AAV-NT-ABE8eWQ, and AAV-CT-ABE) was performed by VectorBuilder.

### Cell culture and transfection

To correct *Rpe65* mutations *in vitro*, mouse embryonic fibroblasts from *rd12* mice were maintained in DMEM containing 10% FBS, 1% penicillin-streptomycin (WELGENE), and 4 mM glutamine (Glutamax-I, Gibco). Then, the 1.0 × 10^5^ cells were electroporated with ABE N-term (335 ng) and ABE C-term (335 ng) using the Neon transfection system (Thermo Fisher Scientific, MPK1025). The electroporation protocol was 1,650 V, 20 ms, 1 pulse.

### Animals

C57BL/6 (stock no. 000664) and *rd12* (stock no. 005379) mice were purchased from the Jackson Laboratory (Bar Harbor, Maine, USA). All animal experiments were approved by the Seoul National University Animal Care and Use Committee and conducted in strict accordance with the guidelines of the Association for Research in Vision and Ophthalmology Statement. Mice were kept under cyclic light (12-on/12-off) with *ad libitum* access to food and water in approved cages.

### Subretinal injection

Mice were anesthetized with an intraperitoneal injection of tiletamine (25 mg/mL)/zolazepam (25 mg/mL) mixture. After anesthetization, mouse eyes were placed in the proper position and pupils were dilated with an eye drop containing phenylephrine hydrochloride (5 mg/mL) and tropicamide (5 mg/mL). The eyelid was opened and protruded to expose the equator for convenient injection. A small hole was punctured at the slight posterior of the limbus using a sterile 30-gauge needle. The 33-gauge blunt needle of microliter syringe was placed through the pre-punctured hole. The needle was inserted into the subretinal space until the point when mild resistance was felt. The solution was injected slowly with low pressure and the retinal bleb was observed under the microscope. Mice received AAV-NT-ABE and ABE-CT-ABE (4.3 × 10^10^ viral genomes for AAV2/2 and 4.3 × 10^10^ viral genomes for AAV2/9 each in 3 μL of PBS) into the subretinal space.

### Immunofluorescence staining

Pups from each group were randomly chosen after 6 weeks of injection, and euthanized by carbon dioxide inhalation. The ocular globe was enucleated and fixed in 4% paraformaldehyde (PFA, P2031; Biosesang, Yongin, KR) for 30 min at room temperature. The cornea and lens were removed, and the retina was dissociated from the retinal pigment RCS complex. The RCS complex was incubated in blocking solution (BP150; Biosolution, Yongin, KR) at room temperature for 2 h and stained with Alexa Fluor 488-conjugated anti-RPE65 antibody (1:100, NB100-355AF488; Novus, Denver, CO, USA) overnight at 4°C. The following day, the stained RCS complex was rinsed three times and incubated in Alexa Fluor 594-conjugated anti-ZO-1 antibody (1:250, 339194; Invitrogen, Carlsbad, CA, USA) at room temperature for 2 h. The samples were counterstained with DAPI (1:1,000; D9542; Sigma-Aldrich, St. Louis, MO, USA) at room temperature for 15 min. The stained RCS complex was placed on a glass slide with the retinal pigment epithelial layer against the glass slide. An adequate amount of mounting solution was added, and a cover slide was placed. Immunostained tissues were observed using a confocal microscope (TCS SP8; Leica, Wetzlar, Germany).

### ERG

After anesthetization and mydriasis were complete, the recording electrode was placed on the corneal surface, and the reference needle electrode was placed subcutaneously on the head. The electrode in the tail served as the ground. Full-field ERG was performed using the electrophysiologic system 3,000 (UTAS E-3000, LKC Technologies Inc., Gaithersburg, MD, USA). Mice were dark adapted for >16 h. Under dark adapted conditions, scotopic responses were recorded using a single dim flash of 0 dB using a notch filter at 60 Hz and a digital bandpass filter of 0.3–500 Hz. The amplitude of the a-wave was measured from the baseline to the lowest negative going voltage, whereas peak b-wave amplitudes were measured from the trough of the a-wave to the highest peak of the positive b-wave. Each group was randomly assigned 8 mice. Among these, 2 and 1 mice in the ABE8e- and ABE8eWQ-treated groups died during the experiment and were excluded from analysis. The ERG waveforms were performed using GraphPad PRISM 7 (GraphPad Software, San Diego, CA, USA).

### OptoMotry response test

A virtual optomotor system (OptoMotry apparatus; CerebralMechanics Inc., Lethbridge, Alberta, Canada) was used to assess visual function. Briefly, the mice were placed on an elevated platform positioned in the middle of an arena created by four inward-facing display monitors. Spatial frequency thresholds were assessed using a video camera to monitor the elicitation of the optokinetic reflex through virtual stimuli projected with sine-wave gratings (100% contrast) on computer monitors. Experimenters were blinded to the treatment and previously recorded thresholds of each animal.

### Targeted DNA and RNA sequencing

The extracted RPE or retina tissue was sonicated with lysis buffer from NucleoSpin RNA Plus Kits (MACHEREY-NAGEL, 740,984.250). Then, one-half of the lysates was purified to prepare genomic DNA using NucleoSpin Tissue Kits (MACHEREY-NAGEL) and the other half was purified to prepare RNA using NucleoSpin RNA Plus Kits (MACHEREY-NAGEL, 740,984.250), according to the manufacturer’s protocol. Purified genomic DNA was amplified using KOD-Multi & Epi (TOYOBO, KME-101) and 1 *μ*L of the PCR product was transferred and further amplified with proper index primers for next-generation sequencing using Illumina Miniseq instrument. The purified RNA was converted into cDNA via reverse transcription using ReverTraAce-α-(TOYOBO, FSK-101), according to the manufacturer’s protocol. The cDNA was amplified with KOD-Multi&Epi (TOYOBO, KME-101) and sequenced by Illumina Miniseq instrument. To obtain the percentage of adenosines edited to inosines, the number of adenosines converted to guanosines was divided by the total number of adenosines in the transcript. All Miniseq results were analyzed using BE-Analyzer (http://www.rgenome.net/be-analyzer/)(Hwang *et al*, 2018).

### AlphaFold-based mutational scan

We used AlphaFold2 to predict the structures of RPE65 and its variants(Jumper *et al*, 2021). The source code is available at https://github.com/deepmind/alphafold. The model with the highest average pLDDT score was used for all predictions. Structural images were generated using PyMOL 2.5.0 (https://github.com/schrodinger/pymol-open-source).

### MD calculation

MD simulations were performed using the GROMACS software package (version 2020.4)(Van der Spoel *et al*, 2005). Simulations were performed using the CHARMM36m force field and TIP3P solvent model to assess structural stability and interaction energy(Huang *et al*, 2017; Jorgensen *et al*, 1983). Each system was equilibrated in a cubic TIP3P water box containing 150 mM Na^+^ and Cl^−^ ions in two steps after steepest descent minimization. For electrostatic interactions, we used the particle mesh Ewald method with a cutoff of 1.2 nm; for van der Waals interactions, the cutoff was 1.2 nm, and a velocity-rescaling thermostat was employed(Bussi *et al*, 2007; Essmann *et al*, 1995). Simulation for 100 ps in an ensemble with a constant volume (NVT) was the first step, and 100 ps constant-pressure (NPT) equilibration was performed with position restraints applied to heavy atoms. Without any restraints, production MD simulations were run for 100 ns. Calculation of interaction energy between the backbone atoms of adjacent domains was performed using the gmx energy module implemented in GROMACS. The structures in the figures were modeled using PyMOL 2.5.0.

### Predictions of protein solubility

To approximate the effect of bystander mutations, we calculated protein solubility at neutral pH using the CamSol web server (http://www-vendruscolo.ch.cam.ac.uk/camsolmethod.html)(Sormanni *et al*, 2015).

### Statistics

All group results are expressed as mean ± SEM, if not stated otherwise. One-way ANOVA and Tukey’s *post hoc* multiple comparison tests were performed for comparison between groups. Statistical analyses were performed using GraphPad PRISM 7.

## Author contributions

J.H.K., and S.B. conceived this project; S.-H.L., Y.K.J. performed the cellular experiments and genotyping analysis; G.-H.H. performed bioinformatic analysis; J.W., H.J., S.J.L performed in vivo experiments; J.W., and D.H.J. analyzed the in vivo results; D.I peformed structural analysis and molecular dynamic simulations; S.-H.L., J.W., D.I., J.H.K., and S.B. wrote the manuscript with the approval of all other authors; W.A.G. provided theoretical analysis and methodology; J.H.K., and S.B. supervised the research.

## Acknowledgements

This work was supported by grants from the National Research Foundation of Korea (NRF) (no. 2022M3A9E4017127 and RS-2023-00260351 to J.H.K.; no. 2021M3A9H3015389, no. 2020M3A9I4036072, RS-2024-00455559 and SRC - NRF2022R1A5A102641311 to S.B.; RS-2023-00274504 to D.I.; RS-2023-00246813 to Y.K.J.), the Korean Fund for Regenerative Medicine (KFRM) RS-2024-00332601 and 21A0202L1-11 to S.B., the Korea Research Institute of Bioscience and Biotechnology (KRIBB) Research Initiative Program (KGM5362111 to J.H.K.), Kun-hee Lee Child Cancer & Rare Disease Project, Republic of Korea (202200004004 to J.H.K.), Seoul National University Hospital Research Grant (18-2023-0010 and 03-2023-3020), and grants from the National Institutes of Health (NIH) R01HL155532 and R35HL150807 to W.A.G.

